# Cooperating yet distinct brain networks engaged during naturalistic paradigms: A meta-analysis of functional MRI results

**DOI:** 10.1101/165951

**Authors:** Katherine L. Bottenhorn, Jessica S. Flannery, Emily R. Boeving, Michael C. Riedel, Simon B. Eickhoff, Matthew T. Sutherland, Angela R. Laird

**Affiliations:** Department of Psychology, Florida International University, Miami, FL; Department of Physics, Florida International University, Miami, FL; Institute of Clinical Neuroscience and Medical Psychology, Heinrich-Heine University, Düsseldorf, Germany; Institute of Neuroscience and Medicine, Research Center Jülich, Jülich, Germany

## Abstract

Cognitive processes do not occur by pure insertion and instead depend on the full complement of co-occurring mental processes, including perceptual and motor functions. As such, there is limited ecological validity to human neuroimaging experiments that use highly controlled tasks to isolate mental processes of interest. However, a growing literature shows how dynamic, interactive tasks have allowed researchers to study cognition as it more naturally occurs. Collective analysis across such neuroimaging experiments may answer broader questions regarding how naturalistic cognition is biologically distributed throughout the brain. We applied an unbiased, data-driven, meta-analytic approach that uses *k*-means clustering to identify core brain networks engaged across the naturalistic functional neuroimaging literature. Functional decoding allowed us to, then, delineate how information is distributed between these networks throughout the execution of dynamical cognition in realistic settings. This analysis revealed six recurrent patterns of brain activation, representing sensory, domain-specific, and attentional neural networks that support the cognitive demands of naturalistic paradigms. Though gaps in the literature remain, these results suggest that naturalistic fMRI paradigms recruit a common set of networks that that allow both separate processing of different streams of information and integration of relevant information to enable flexible cognition and complex behavior.

## Introduction

Across the life sciences, researchers often seek a balance between ecological validity and careful laboratory control when making experimental design decisions. This entails weighing the value of creating realistic stimuli representative of real-world, interactive experiences versus artificial, reductionist stimuli facilitating precise assessment of ‘isolated’ mental process of interest via cognitive subtraction. Cognitive subtraction assumes that a single added cognitive process does not alter the other, co-occurring processes, both neutrally and cognitively. As such, task-based fMRI has traditionally utilized precisely controlled tasks to study the neurobiological substrates of cognition. However, cognition does not occur by pure insertion; the functioning of any cognitive process is not wholly independent from other co-occurring processes (Friston et al., 1996). Instead, cognition is highly interactive, encompassing measurable changes in neural activity that are dependent on the full amalgamation of relevant social, cognitive, perceptual, and motor processes. Thus, it is perhaps unreasonable to expect findings from a highly restricted assessment of a psychological construct in the scanner to fully generalize to real-world behaviors and settings.

With advances in technology and a desire to study cognition with greater ecological validity, increasing numbers of studies are utilizing realistic, interactive, and rich stimuli in more ecologically valid experimental designs that fit within the scanner’s confines (Hasson and Honey, 2012; Maguire, 2012; Wang et al., 2016). “Naturalistic” paradigms employ dynamic and complex stimuli (Fehr et al., 2014; Kauttonen et al., 2015; Burunat et al., 2014), in terms of multimodal demands (Lahnskoski et al., 2012; Maguire et al., 2012; Nardo et al., 2014; Dick et al., 2014; Reed et al., 2014; Bishop et al., 2014), or in relation to the length of the stimulus presentation (Maguire et al., 2012; Cong et al., 2014). Specifically, the use of video games, film clips, and virtual reality, among others, has brought a new dimension to cognitive neuroimaging experiments permitting researchers to study brain activity as participants engage in tasks that more closely represent real-life demands on attention and multimodal sensory integration. Appreciation of such attention and integration processes necessitates more complex stimuli than simple static images presented on a screen. For example, researchers have studied spatial navigation with virtual reality environments as complex as the city of London (Spiers and Maguire, 2006) and as classic as a virtual radial arm maze (Marsh et al., 2010). Similarly, social cognition has been probed with displays of human social interactions from a dramatic, social television drama (Spunt and Lieberman, 2012) to clips of facial expressions with little context (Li et al., 2015).

Everyday activities, such as navigation or social observation, involve the integration of processes associated with object recognition, speech comprehension, motor control, and spatial orienting, which all require the interpretation of dynamic signals often from more than one sensory modality (e.g. audiovisual film watching or visuotactile image tracing) and necessitate different attentional demands compared to the simplistic stimuli used in traditional fMRI experiments (Giard and Peronnet, 1999; McGurk and MacDonald, 1976; Sailer et al., 2000; Spence, 2010). Recently, this trend has produced open-source efforts such as studyforrest, a freely-available dataset of MRI scans, eye-tracking, and extensive annotations, using the movie Forrest Gump as a rich, multimodal stimulus (studyforrest.org; Hanke et al., 2016; 2015; 2014). Although studies of participants freely viewing films or navigating virtual environments have been used since the early days of fMRI, the naturalistic studies represent a small portion of the overall task-based fMRI literature (Beauregard et al., 2001; Burgess et al., 2001; Maguire, 2012). Despite offering advantages, the growing body of naturalistic fMRI research has yet to be quantitatively assessed, and little is known of how the neural bases of these tasks support complex information processing and behavioral demands.

Here, we applied an unbiased, data-driven, meta-analytic approach to quantitatively explore and classify knowledge embedded in the naturalistic fMRI literature. Using an approach developed by Laird et al. (2015), we capitalized on the wealth and flexibility of published naturalistic paradigms and investigated recurrent patterns of brain activation reported across a wide variety of tasks and behaviors of interest. This method is based on the premise that functionally similar tasks engage spatially similar patterns of brain activity and that, by clustering activation patterns from experimental contrasts, similar of experimental paradigms can be identified. Naturalistic paradigms are uniquely rich here, due to the multitude of component processes contributing to realistic behavior that can be illuminated by modeling strategies in data analysis. To this end, we extracted relevant information about the stimuli and task demands of these paradigms and assessed motifs in the arrangement of this information, with respect the data-driven clustering analysis, to determine which paradigm aspects elicited activation patterns that subserve common and dissociable cognitive processes. Although naturalistic paradigms vary greatly and are designed to probe a wide range of psychological constructs and behaviors, we hypothesized that complex, multisensory processing are associated with a set of core neural networks engaged by similar content domains and task demands. The objectives of this study were to first elucidate core brain networks engaged by the myriad processes that underlie behavior during naturalistic fMRI paradigms and, then to characterize how information processing is potentially distributed between these networks to facilitate complex behaviors in realistic settings.

## Methods

### Naturalistic fMRI Paradigms

Here, “naturalistic” paradigms were operationally defined as tasks employing any stimulus which demanded continuous, real-time integration of dynamic streams of information. This definition excludes any paradigms based on still-frame stimulus presentation, which intrinsically impose static constraints that are rarely present in the world and, thus, limit ecological their validity. Importantly, a key distinction of naturalistic tasks is that stimuli are continuously presented across the duration of the task, while other tasks in the literature rely on repeated trials of stimuli. As real-world behavior contextually involves all sensory modalities, we included naturalistic tasks in which such stimuli were presented via the visual, auditory, or tactile modalities or any combination thereof. Visual naturalistic tasks require either a real-time interaction with visual stimuli, in the case of video games and virtual reality, or the continuous integration of real-time information, such as during film viewing. Auditory tasks, including the perception of music and spoken stories, similarly require the continuous integration of, and often interaction with, real-time information. Our operational definition also included tactile naturalistic paradigms, which involve the manipulation and recognition of physical objects. During these tactile tasks, participants gather and integrate sensory information to create a mental representation of the object and, if necessary, form an appropriate behavioral response. Lastly, we note the inclusion of multisensory tasks. As in life, many naturalistic experiments simultaneously present auditory, visual, and tactile information, and such tasks demand the real-time integration of information from multiple sensory modalities.

### Literature Search, Filtering, and Annotation

An extensive literature search was performed to amass a corpus of naturalistic fMRI studies that were published since the emergence of fMRI in 1992. To identify published naturalistic fMRI studies, PubMed searches were carried out by focusing on stimulus types common to naturalistic research (e.g., video games, film, virtual reality). The first search string, performed on January 13, 2016, used the following string to identify relevant studies by their titles and abstracts: ((“naturalistic”[Title/Abstract] OR “real-world”[Title/Abstract] OR “ecologically valid”[Title/Abstract] OR “true-to-life”[Title/Abstract] OR “realistic”[Title/Abstract] OR “video game”[Title/Abstract] OR “film”[Title/Abstract] OR “movie”[Title/Abstract] OR “virtual reality”[Title/Abstract]) AND (“fMRI”[Title/Abstract] OR “functional magnetic resonance imaging”[Title/Abstract]) AND (“Humans”[MeSH])). This search yielded 679 studies (January 2016), some of which utilized stimulus types that we had not included in our initial query, including music, speech, and tactile objects. To identify any studies using these tasks that may not have been returned by initial query, a second search was performed on January 20, 2016 using the string [(“music”[Title/Abstract] OR “speech”[Title/Abstract] OR “spoken”[Title/Abstract] OR “tactile object”[Title/Abstract]) AND (“naturalistic”[Title/Abstract] OR “real-world”[Title/Abstract] OR “ecologically valid”[Title/Abstract] OR “true-to-life”[Title/Abstract] OR “realistic”[Title/Abstract]) AND (“fMRI”[Title/Abstract] OR “functional magnetic resonance imaging”[Title/Abstract]) AND “Humans”[MeSH]]. This secondary search returned 48 studies, some of which were included in the results of the first search. The two sets of search results were pooled to identify 754 unique studies, which were then reviewed and filtered to identify studies utilizing naturalistic paradigms as defined above.

Each of 754 candidate studies was first screened and then reviewed according to the following exclusion criteria (Figure 1; Moher, Liberati, Tetzlaff, Altman, & Altman, 2009). The screening process examined the Abstracts and Methods of each paper to exclude non-naturalistic tasks in which static, timed blocks of stimuli were presented with a well-defined window for participant response. In this step, we also excluded studies that assessed training or learning across multiple trials or across some period of practice (e.g., pre vs. post contrasts), as our focus was on neural underpinnings of the tasks themselves and not training-induced changes thereof. In determining eligibility of each paper, studies of participants under the age of 18 or of participants with any history of neurological or psychiatric diagnosis were excluded. After this study-level examination, we then inspected each reported experimental contrast within each paper. In this context, “experiment” represents each statistical parametric image presented, as the result of some functional image data analysis, such as contrasting experimental conditions (Fox et al., 2005). Experiments from analyses that used an *a priori* region(s) of interest to investigate activation or functional connectivity were omitted permitting identification of whole-brain neural networks. We also excluded contrasts modeling ANOVA interaction-specific activations, due to the inherent complexity of such effects. In this step, any studies/contrasts that did not meet the minimum requirements for coordinate-based meta-analysis, reporting the brain activation locations in a three-dimensional, standardized coordinate space, were discarded.

**Figure 1.**
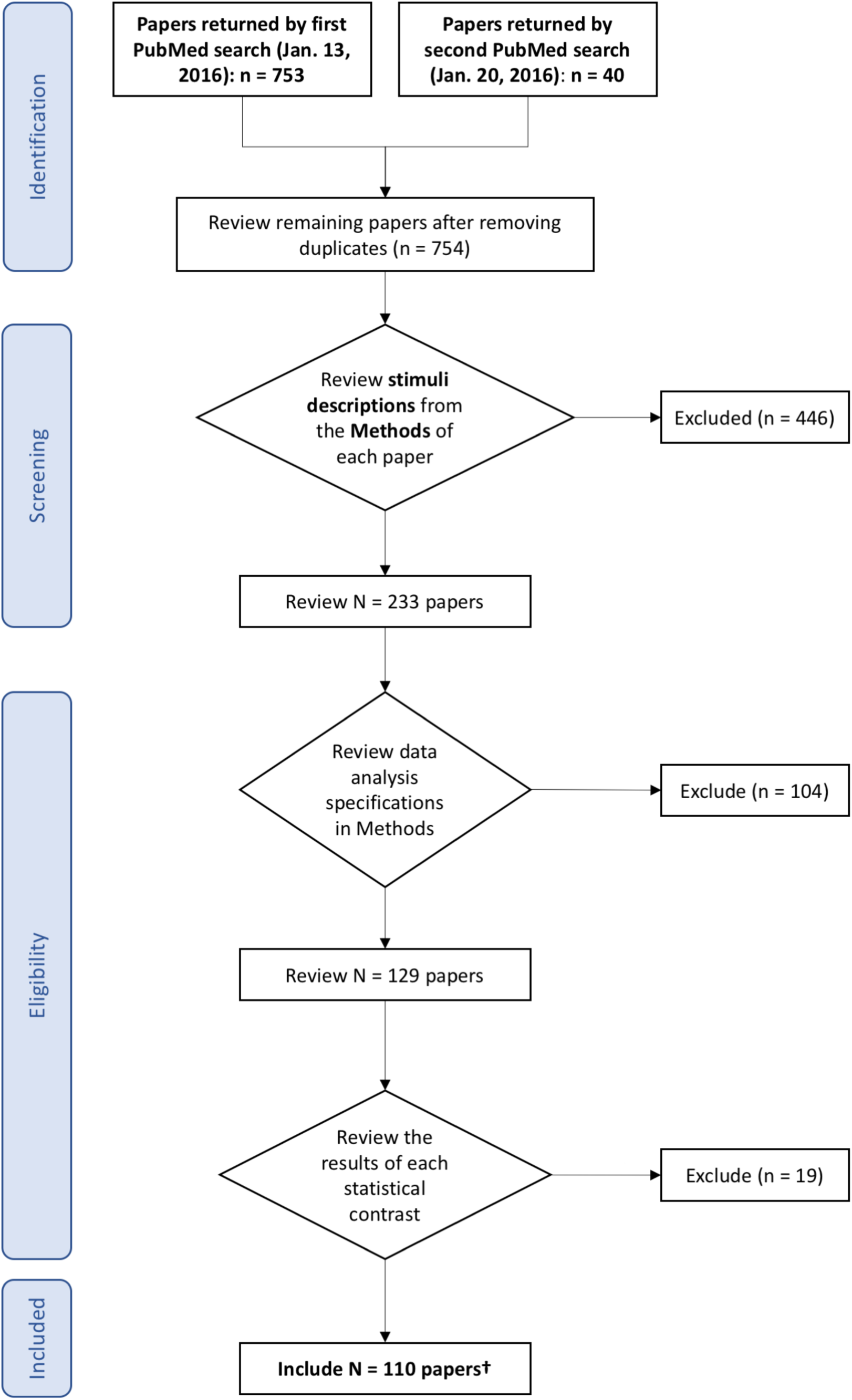
PRISMA flow chart of inclusion and exclusion criteria. Each of the experiments returned by the PubMed queries were screened according to this schematic.

During inspection of each contrast, one study associate (KLB) manually annotated each experiment with terms that described the experimental design with respect to stimulus type utilized, sensory modality engaged, and the task nature. These terms described the salient aspects of the stimuli and behaviors associated with each individual experimental contrast from the corpus of naturalistic paradigms, annotating the particular aspects of the tasks highlighted by each modeled experimental contrast, and not the intended psychological construct interrogated by the original report. These manual annotations were then independently reviewed and confirmed by a second study associate (JSF) to assure consistency and accuracy. Any disagreements or inconsistencies between KLB and JSF were resolved following a final conversation between the two associates.

### Experimental Design and Statistical Analysis

#### Modeled Activation Maps

Following the identification of relevant papers and experiments/contrasts, reported brain activation coordinates were extracted. All Talairach atlas-based coordinates (Talairach and Tournoux, 1988) were converted to Montreal Neurological Institute (MNI) space (Collins et al., 1994; Evans et al., 1993) using the tal2icbm transformation (Lancaster et al., 2007; Laird et al., 2010). Probabilistic modeled activation (MA) maps were created from the foci reported in each individual contrast by modeling a spherical Gaussian blur around each focus with FWHM determined by the number of subjects in each experiment in order to represent the uncertainty induced by the inherent variability from individual differences and between-lab differences (Eickhoff et al., 2009). These MA maps were concatenated into an array of *n* experiments by *p* voxels, which was then analyzed for pairwise correlations that reflected the degree of spatial similarity between the MA maps from each of the *n* experiment and those of every other experiment. The resultant *n* X *n* correlation matrix represented the similarity of spatial topography of MA maps between every possible pair of experiments.

#### K-Means Clustering Analysis

Individual naturalistic experiments (*n* MA maps) were then classified into *K* groups based on their spatial topography similarities. The *k*-means clustering procedure was performed in Matlab (Mathworks, R2013b for Linux), which grouped experiments by pairwise similarity, calculating correlation distance by one minus the correlation between MA maps (from the aforementioned correlation matrix) and finding the “best” grouping by minimizing the sum of correlation distances within each cluster (code available at https://github.com/62442katieb/meta-analytic-kmeans). This approach begins by choosing *K* arbitrary maps as representative centroids for each of *K* clusters and assigning experiments to each cluster based on the closest (most similar) centroid. This process continued iteratively until a stable solution was reached.

Solutions were investigated for a range of *K* = 2 – 10 clusters. Once the clustering analysis was complete for all *K*, we compared each solution to the neighboring solutions and assessed for improvement across parcellation schemes using four metrics describing cluster separation and stability (Bzdok et al., 2015; Eickhoff et al., 2016a). This allowed us to objectively select the number of clusters that most optimally divided the data set. The first metric, *average cluster silhouette* across clustering solutions, assessed the separation between clusters and described whether clusters were distinct or overlapping. A higher silhouette value indicates that greater separation is ideal and that each experiment fits well into its cluster, with lower misclassification likelihood of fringe experiments into neighboring clusters. Stability is indicated by a relatively minimal change in silhouette from one solution (*K*) to the next (*K* + 1), indicated by the positive derivative of the silhouette score closest to zero, with greatest stability evidenced by the smallest change between two points. Second, we considered the *consistency of experiment assignment* by comparing the ratio of the minimum number of experiments consistently assigned to a cluster relative to the mean number of experiments consistently assigned to that cluster. In this case, only ratios above 0.5, in which at least half of the experiments were consistently assigned, were considered viable solutions. Third, the *variation of information* was quantified, which compared the entropy of clusters with the mutual information shared between them for each solution *K* and its *K* – 1 and *K* + 1 neighbors. A large decrease in variation of information from *K* – 1 to *K* and increase from *K* to *K* + 1, a local minimum in the plot of variation of information across *K*, indicated a decrease in overlap between solutions and, thus, stability of solution *K*. In this case, “large” is defined, too, in relative terms, with the largest decrease indicating greatest stability of the solutions considered. Finally, we computed a *hierarchy index* for each solution, which assessed how clusters split from the *K* – 1 to *K* solution to form the additional cluster. A lower hierarchy index indicated that clusters present in *K* stemmed from fewer of the clusters present in *K* – 1, another indication of stability in groupings demonstrated by a local minimum across values of *K*. An optimal clustering solution is one that demonstrated minimal overlap between clusters (i.e., high silhouette value), while exhibiting relative stability in comparison with the previous and next solutions (i.e., consistency > 0.5, a local minimum in variation of information, and lower hierarchy index than previous).

#### Meta-Analytic Groupings

From the identified optimal clustering solution, we probed the underlying neural topography associated with each of the *K* groups of experiments (Laird et al., 2015). To this end, the ALE meta-analysis algorithm (Turkeltaub et al., 2002; Laird et al., 2005) was applied to generate a map of convergent activation for each grouping of experiments with similar topography. The ALE algorithm includes a weighting of the number of subjects when computing these maps of convergent activation and accounts for uncertainty associated with individual, template, and registration differences between and across experiments (Eickhoff et al., 2009; Turkeltaub et al., 2012). The union of these probability distributions was used to calculate ALE scores, a quantitative assessment of convergence between brain activation across different experiments, which was compared against 1000 permutations of a null distribution of random spatial arrangements (Eickhoff et al., 2012). These ALE values for each meta-analytic grouping of experiments were thresholded at *P* < 0.01 (cluster-level corrected for family-wise error) with a voxel-level, cluster-forming threshold of *P* < 0.001 (Eickhoff et al., 2016b; Woo et al., 2014). The resultant ALE maps thus reflected the convergent activation patterns within each of the *K* clusters. The experimental *K* clusters are hereafter referred to as meta-analytic groupings (MAGs), representing meta-analytic groups of experiments demonstrating similar activation patterns.

### Functional Decoding

Once we elucidated convergent activation patterns within MAGs, we sought to gain insight into what aspects of the naturalistic paradigms were most frequently associated with each MAG via functional decoding. Functional decoding is a quantitative, data-driven method by which researchers can infer which mental processes are related to activation in a specific brain region (or set of brain regions) across published fMRI studies. We chose to use two complementary functional decoding approaches, one based on our study-specific, subjective manual annotations mentioned above, and another based on the objective, automated annotations provided by the Neurosynth database for over 11,000 functional neuroimaging studies (Yarkoni et al., 2011; Neurosynth.org). First, the manually annotated terms associated with each experiment were grouped into the MAGs identified above and were assessed by frequency of occurrence in each MAG. The distribution of stimulus modality, stimulus type, and salient terms across MAGs allowed us to evaluate the relationship between activation patterns and the aspects of naturalistic paradigms that elicited them. Second, we included an automated, data-driven annotation method using Neurosynth, which includes automatically extracted terms that occur at a high frequency in the abstract of each archived study. To functionally decode our MAGs, we compared the MAGs’ activation patterns with those reported across published neuroimaging papers in the Neurosynth database. To this end, we uploaded each ALE map to NeuroVault, a web-based repository for 3D statistical neuroimaging maps that directly interfaces with Neurosynth (Gorgolewski et al., 2015; NeuroVault.org,). NeuroVault enables “functional decoding” by correlating unthresholded uploaded maps with term-specific meta-analytic maps extracted from Neurosynth’s database of published functional neuroimaging studies. The Neurosynth functional decoding results were exported as a set of terms and correlation values representing how well the spatial distribution of activation associated with each term in the database matched the activation pattern of the uploaded map.

Both sets of terms (i.e., obtained via manual and automated approaches) were evaluated to assess the specific aspects of naturalistic paradigms associated with each MAG. The Neurosynth terms representing broad behavioral aspects across fMRI studies that elicit similar brain activation profiles provides both an unbiased description of the experiments engaging each MAG, as well as a comparison of our corpus of studies with the broader literature. On the other hand, manual annotation provides more concise, accurate description of the paradigms, though it is predisposed to the subjective bias of human annotation. The results of this two-pronged functional decoding approach were designed to describe the processes that engage brain networks similar to each MAG and how these processes may be similar or different in naturalistic fMRI studies compared to the broader functional neuroimaging literature. The distribution of stimulus modalities and types across MAGs was assessed, too. Together, the functional decoding results and distributions of different stimuli were interpreted to provide insight into how information processing is functionally segregated across cooperating neural systems during naturalistic tasks.

## Results

The literature search yielded a combined set of 110 studies that reported coordinates of brain activation from naturalistic fMRI tasks among healthy adults (Figure 1, PubMed IDs available in Supplementary Table 1). The final data set included activation foci from 376 experimental contrasts (*N* = 1,817 subjects) derived from tasks using a variety of stimulus types and sensory modalities. Across our corpus of naturalistic fMRI experiments, approximately 55% assessed a single stimulus modality, including 40% visual stimuli, 13% auditory, and 1% tactile.

**Table 1.** Distribution of stimulus modalities across the naturalistic corpus. Paradigms engaged auditory, visual, and tactile sensory modalities, both separately and in combination.

| Stimulus Modality | Number of Experiments |
| --- | --- |
| Auditory | 50 (13%) |
| Audiovisual | 154 (41%) |
| Visual | 150 (40%) |
| Visual + tactile (pain) | 9 (2%) |
| Visual + tactile | 5 (1%) |
| Tactile | 4 (1%) |

Conversely, 45% of experiments utilized multisensory stimuli, including 41% that employed audiovisual stimuli, 2% in which a visual stimulus was paired with painful, tactile stimuli, and 1% pairing visual and non-painful tactile stimuli (Table 1). Of the visual experiments, 69% involved a motor response, as did 25% of the audiovisual experiments, ranging from a button press to joystick and object manipulation. The stimulus types most frequently used across the included experiments were films (45%), virtual reality (32%), speech (9%), and music (6%) (Table 2).

**Table 2.** Distribution of stimulus types across the naturalistic corpus. Within each stimulus modality, multiple types of experimental stimuli were included across the data set.

| Stimulus Type | Number of Experiments |
| --- | --- |
| Film | 169 (45%) |
| Virtual Reality | 121 (32%) |
| Speech | 32 (9%) |
| Music | 21 (6%) |
| Video Game | 13 (4%) |
| 3D image | 6 (2%) |
| Tactile | 6 (2%) |
| Picture | 4 (1%) |
| Sounds | 1 (<1%) |

## K-Means Clustering Solutions

MA maps were created for each contrast, and then clustered to identify groups with similar activation topographies. For completeness, the *k*-means clustering solutions for *K* = 2 – 10 clusters were quantitatively evaluated across four metrics to identify an optimal solution (Figure 2). When considering the average silhouette metric (Fig. 2A), values generally increased as *K* increased and the *smallest increase* was observed between *K* = 6 to *K* = 7, indicating little additional separation between clusters gained by moving from 6 to 7 clusters. With respect to the consistency of assigned experiments metric (Fig. 2B), each of the solutions *K* = 2 – 10 met the stability requirement whereby the minimum number of experiments included in any iteration of the solution was at least 50% of the mean number of experiments included across iterations. The variation of information metric (Fig. 2C), suggested the stability of a 6-cluster solutions as parameter value *decreases* were observed when moving from *K* = 5 to *K* = 6, combined with parameter *increases* when moving from *K* = 6 to *K* = 7, indicating that a 6-cluster solution demonstrates relative stability. The hierarchy index metric (Fig.2D) further corroborated a 6-cluster solution, as a local minimum as observed at *K* = 6. Due to agreement across these metrics, we chose to proceed with the *K* = 6 solution.

**Figure 2.**
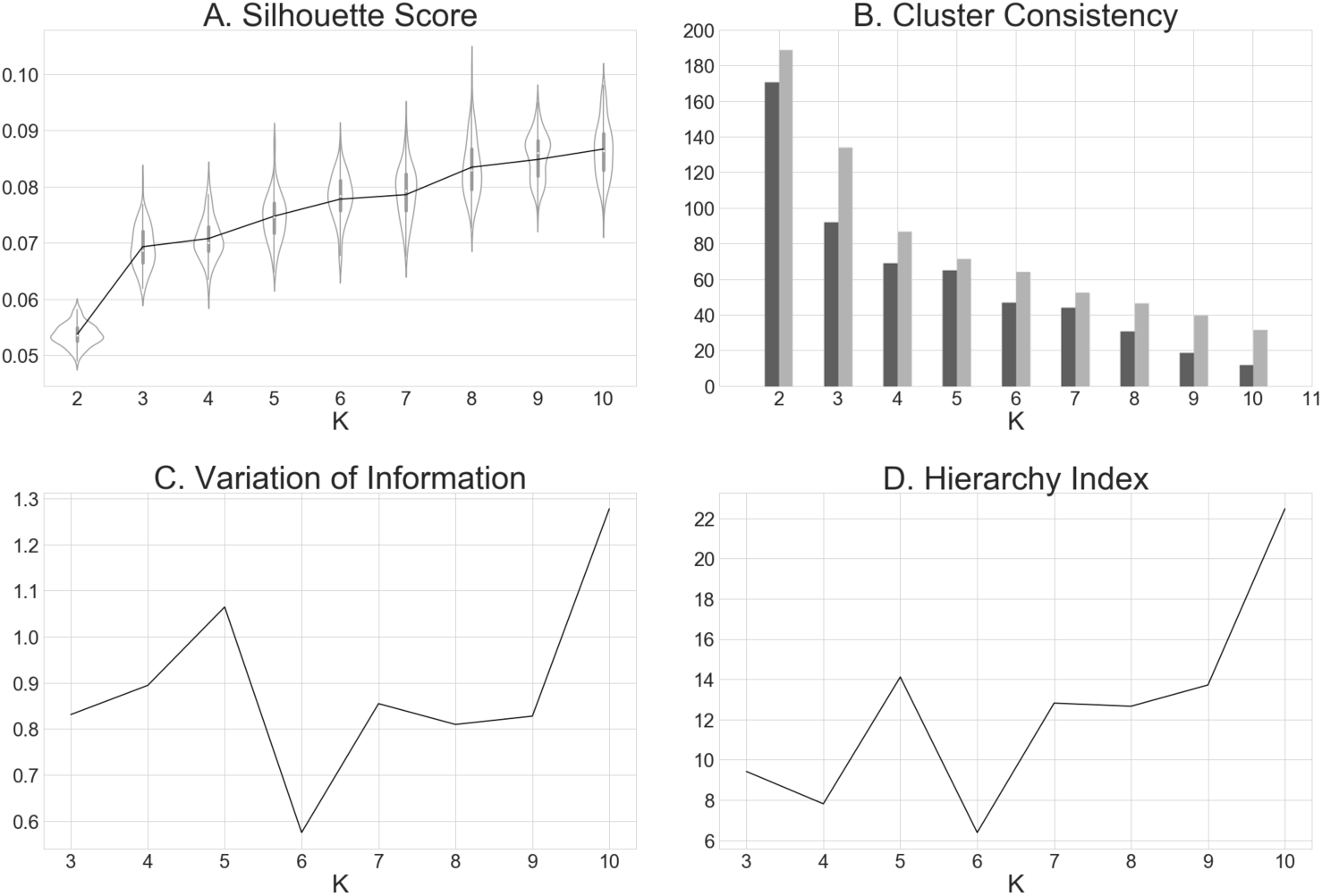
Metrics computed for *K* = 2 – 10 clustering solutions. (A) The average cluster silhouette for each solution *K* from 2 to 10 clusters, showing the distribution of average silhouette values at each value of *K*, resampled 100 times leaving one random experiment out each time. (B) Consistency in experiments assignment to clusters, plotting the minimum consistently assigned clusters next to the mean of consistently assigned clusters. (C) The change in variation of information, a distance metric, from the *K* – 1 to *K* and from *K* to *K* + 1. (D) The hierarchy index for each of *K* clustering solutions, which provides information about how clusters in the *K* solution stemmed from clusters in the *K* – 1 solution.

## Meta-Analytic Groupings

The optimal clustering solution yielded six meta-analytic groupings (MAGs) of experiments in our corpus, suggesting similarities in brain activation across this sample of the naturalistic literature coalesce into six distinct patterns. The number of experiments that were clustered into each MAG ranged from 50 to 83 experiments (*mean* = 62.67; SD= 12.46). ALE maps of the six MAGs were generated and demonstrated little overlap in activation patterns, suggesting distinct patterns of recurrent activation across our set of naturalistic experiments (Figure 3, Supplementary Table 2). Whereas some of the MAGs exhibited focal patterns of convergent activation, restricted to a single or neighboring gyri (e.g., MAG 1 and 5), others presented with distributed convergence across multiple lobes (e.g., MAG 2 and 6). Most of the resulting MAGs were restricted to cortical activation patterns, although MAG 3 exhibited convergent activation in subcortical and brainstem regions (results available on NeuroVault at https://neurovault.org/collections/3179/).

**Figure 3.**
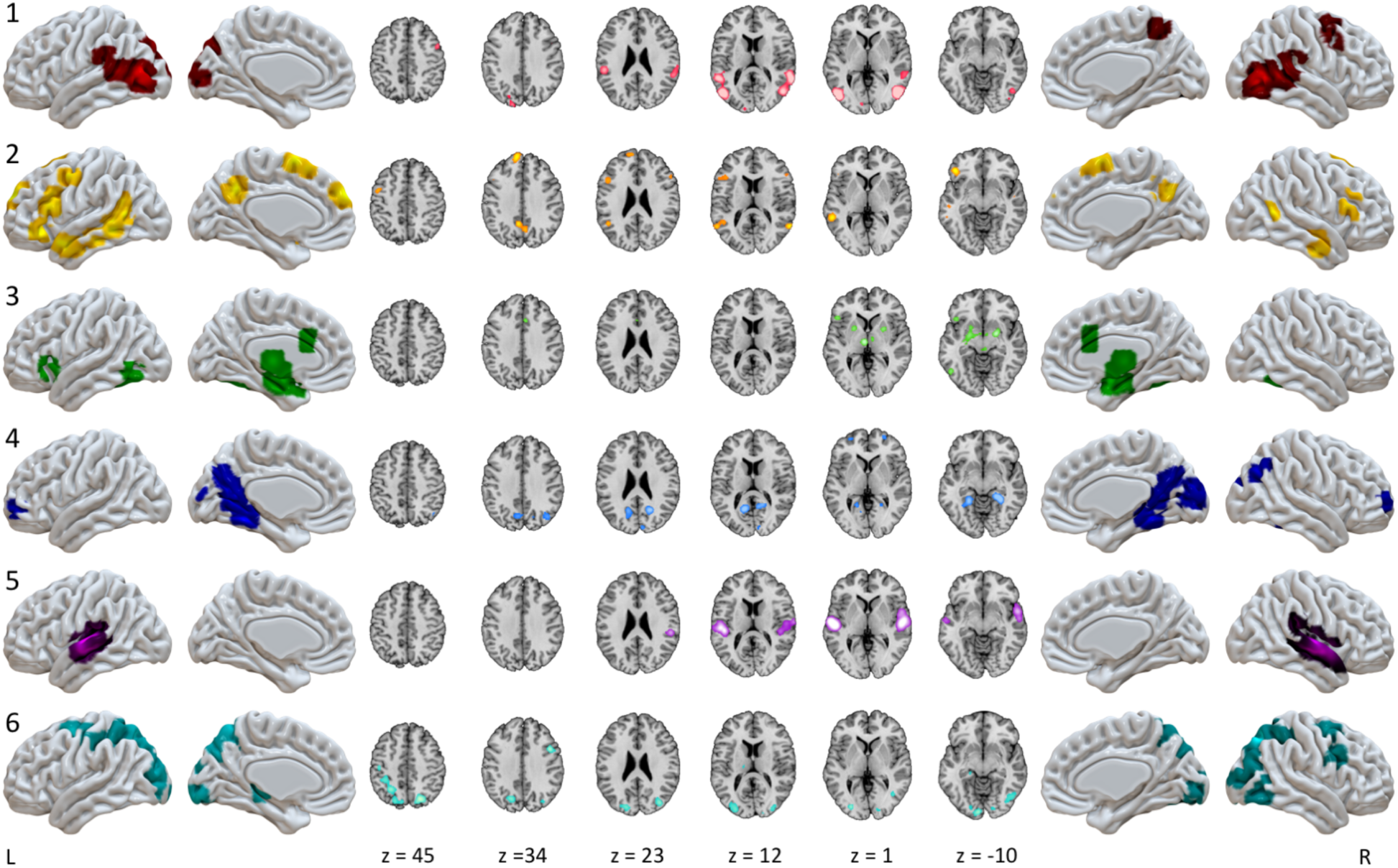
Convergent activation patterns of MAGs from the naturalistic corpus. ALE meta-analysis of experiments in each MAG yielded six patterns of convergent activation.

***MAG 1*** included convergent activation in the bilateral posterior temporal areas, including portions of the inferior, middle, and superior temporal gyri, extending into the inferior parietal lobule and into the middle occipital gyrus, as well as in the left supramarginal gyrus, right precentral and middle frontal gyri, and in the bilateral precuneus. ***MAG 2*** exhibited convergence in left inferior frontal gyrus, left precentral gyrus, anterior and posterior aspects of the middle temporal gyrus, precuneus, in addition to both the left and right superior frontal gyri. ***MAG 3*** demonstrated a largely symmetric convergence pattern across multiple subcortical structures including bilateral amygdalae, putamen, thalamus, parahippocampal gyrus, and periaqueductal gray, with cortical clusters observed in the left inferior frontal sulcus and inferior frontal gyrus, bilateral anterior cingulate cortex, and bilateral fusiform gyri. ***MAG 4*** exhibited convergent activation in bilateral medial temporal lobes, parahippocampal regions, bilateral precuneus, retrospenial posterior cingulate cortex, occipital regions including the lingual gyrus, right calcarine sulcus, and cuneus, in addition to a small, bilateral portion of the middle frontal gyri. ***MAG 5*** showed convergence in the bilateral superior temporal gyri. ***MAG 6*** demonstrated convergence in the bilateral superior frontal sulci, intraparietal sulci, and superior parietal lobules as well as convergence in higher-order visual processing areas in the middle occipital and lingual gyri.

## Stimulus Distribution Across MAGs

Each stimulus modality was represented in multiple MAGs, but modalities were not evenly distributed across MAGs (Figure 4A). Experiments utilizing audiovisual tasks were somewhat uniformly distributed across the MAGs, with a slightly higher proportion of audiovisual tasks in MAGs 1, 3, and 5. In contrast, more than half of the experiments using auditory tasks were grouped into MAGs 2 and 6. Notably, more experiments based on auditory and audiovisual stimuli were clustered into MAG 5 than any other MAG. Experiments in which participants experienced physical pain were not present in MAGs 1, 5, and 6, but distributed nearly evenly among MAGs 2 through 4, with a slightly higher portion in MAG 3. More than half of experiments that used tactile stimuli were grouped into MAG 5 and 6. Visual experiments were more evenly distributed across clusters, though there was a markedly smaller proportion in MAG 5 than any other MAG. One stimulus type, “Sounds”, was represented only once across the corpus and was, thus, excluded from Figure 4. The complete distribution of stimulus modalities across MAGs is provided in Supplementary Table 3.

**Figure 4.**
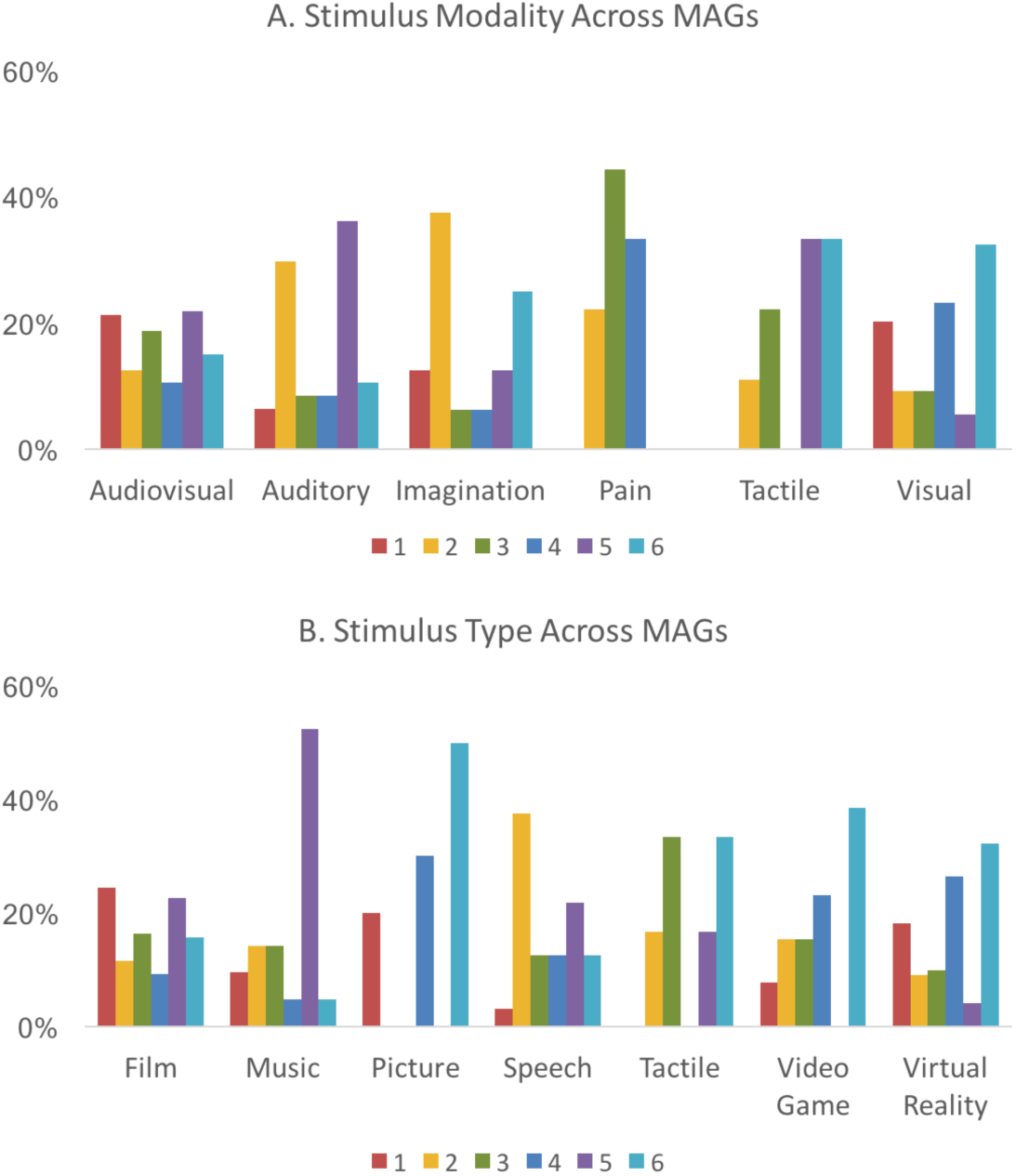
Distribution of stimulus modalities and types across MAGs. (A) The presence of each sensory modality across the corpus that is associated with each MAG. (B) The proportion of each stimulus type present within the corpus that is associated with each MAG. These percentages represent the proportion modality or stimulus type present in each MAG, compared to the total count of that modality or stimulus type across all MAGs.

As with stimulus modality, most stimulus types showed unequal, but not necessarily selective, distribution across MAGs (Figure 4B). Film-based experiments were uniformly distributed across MAGs and tasks utilizing spoken stimuli were more frequently grouped into MAGs 2 and 5. Again, auditory stimuli were highly associated with MAG 5, as more than 50% of music experiments and 20% of speech experiments were clustered into MAG 5. Experiments that required subjects to play video games were most often grouped into MAGs 4, and 6. Experimental contrasts which included a condition in which participants received tactile stimulation or manipulated tactile objects, were most prevalent in MAGs 3 and 6. A detailed distribution of stimulus types across MAGs is shown in Supplementary Table 4.

## Functional Decoding

Two approaches for functionally decoding each MAG, manual and automated annotations, were performed to develop a functional interpretation of each MAGs’ association with aspects of naturalistic paradigms.

### Manual Annotations

Our manual annotations utilized a list of 26 corpus-specific metadata terms, which captured salient features of the naturalistic design, rather than the psychological constructs assumed to be involved. Table 4 displays each of these terms and their frequency of occurrence across MAGs and across the entire corpus (Column = “Total”), highlighting which terms described the largest number of experiments (e.g., “*navigation*”, “*visual features*”, “*emotional film*”, “*attention*”), as well as those that accounted for a minimal number of experiments (e.g., “*violence*”, “*tactile*”, “*pain*”). Values in Table 4 indicate the percent of experiments labeled with each term, or the base-rate of each term throughout the data set, keeping in mind that each experiment was labeled with only one or two terms. Once the experiments were clustered into six MAGs, we evaluated the relative contributions of each term per MAG, controlling for base-rate by dividing each term’s per-MAG count by that term’s total count across the corpus (Table 4). We assessed, too, the ability of each term to predict whether an experiment labeled with that term will be clustered into each MAG, (P(MAG|term) or “forward inference”, and the ability of belongingness to each MAG to predict whether an experiment will be labeled with a particular term, (P(term|MAG)) or “reverse inference”. These outcomes provide the association of each term with each MAG (Table 4). Some of the terms in the manual annotation analysis corresponded to stimulus types in Figure 4B (e.g., per-MAG distribution for “*music*” and “*video game*”). However, many of the manually derived terms highlighted experimental aspects that reflect the unique and salient features of the naturalistic corpus (e.g., “*anthropomorphic*”, “*violence*”) and are not included in standard neuroimaging paradigm ontologies such as BrainMap (Fox et al., 2005) or CogPO (Turner and Laird, 2012).

**Table 4.** Manual functional decoding results across meta-analytic groupings. The relative contributions of each manually-derived metadata term (e.g., term frequencies) were computed for all MAGs, controlling for the base-rate by dividing each term’s per-MAG count by that term’s total count across the corpus. Base-rates are provided as the total count for each term.

| Term | Total |  | Frequency per MAG |  |  |  |  |  |  |  |  |  |  |  |
| --- | --- | --- | --- | --- | --- | --- | --- | --- | --- | --- | --- | --- | --- | --- |
|  |  |  | MAG 1 |  | MAG 2 |  | MAG 3 |  | MAG 4 |  | MAG 5 |  | MAG 6 |  |
| anthropomorphic | 21 | 3% | 10† | 48% | 0 | 0% | 2 | 10% | 2 | 10% | 4 | 19% | 3 | 14% |
| attention | 50 | 7% | 18*† | 36% | 3 | 6% | 2 | 4% | 8 | 16% | 10* | 20% | 9 | 18% |
| auditory features | 17 | 3% | 0 | 0% | 1 | 6% | 1 | 6% | 2 | 12% | 12*† | 71% | 1 | 6% |
| congruence | 22 | 3% | 7 | 32% | 4 | 18% | 0 | 0% | 2 | 9% | 3 | 14% | 6 | 27% |
| emotional film | 61 | 9% | 17* | 28% | 8 | 13% | 17*† | 28% | 4 | 7% | 11* | 18% | 4† | 7% |
| encoding | 24 | 4% | 1 | 4% | 3 | 13% | 1 | 4% | 6 | 25% | 0 | 0% | 13*† | 54% |
| erotic | 15 | 2% | 1 | 7% | 0 | 0% | 8† | 53% | 1 | 7% | 0 | 0% | 5 | 33% |
| faces | 21 | 3% | 5 | 24% | 2 | 10% | 2 | 10% | 2 | 10% | 8† | 38% | 2 | 10% |
| imagination | 23 | 3% | 4 | 17% | 6 | 26% | 2 | 9% | 2 | 9% | 4 | 17% | 5 | 22% |
| inference | 11 | 2% | 4 | 36% | 6† | 55% | 0 | 0% | 1 | 9% | 0 | 0% | 0 | 0% |
| language | 47 | 7% | 9 | 19% | 11* | 23% | 3 | 6% | 4 | 9% | 14*† | 30% | 6 | 13% |
| movement | 14 | 2% | 4 | 29% | 0 | 0% | 1 | 7% | 2 | 14% | 2 | 14% | 5 | 36% |
| music | 21 | 3% | 2 | 10% | 3 | 14% | 3 | 14% | 1 | 5% | 11*† | 52% | 1 | 5% |
| narrative | 30 | 4% | 5 | 17% | 5 | 17% | 1 | 3% | 4 | 13% | 11*† | 37% | 4 | 13% |
| navigation | 81 | 12% | 8* | 10% | 7 | 9% | 10* | 12% | 26*† | 32% | 2 | 2% | 28* | 35% |
| negative valence | 27 | 4% | 8 | 30% | 3 | 11% | 9 | 33% | 1 | 4% | 4 | 15% | 2 | 7% |
| pain | 9 | 1% | 0 | 0% | 2 | 22% | 4† | 44% | 3 | 33% | 0 | 0% | 0 | 0% |
| positive valence | 11 | 2% | 2 | 18% | 4 | 36% | 2 | 18% | 2 | 18% | 1 | 9% | 0 | 0% |
| recognition | 12 | 2% | 0 | 0% | 4 | 33% | 2 | 17% | 1 | 8% | 1 | 8% | 4 | 33% |
| retrieval | 23 | 3% | 1 | 4% | 5 | 22% | 2 | 9% | 4 | 17% | 1 | 4% | 10 | 43% |
| social | 26 | 4% | 9 | 35% | 8 | 31% | 2 | 8% | 3 | 12% | 1 | 4% | 3 | 12% |
| spatial memory | 10 | 1% | 0 | 0% | 2 | 20% | 0 | 0% | 7† | 70% | 0 | 0% | 1 | 10% |
| tactile | 9 | 1% | 0 | 0% | 1 | 11% | 1 | 11% | 0 | 0% | 3 | 33% | 4 | 44% |
| video game | 15 | 2% | 1 | 7% | 2 | 13% | 2 | 13% | 4 | 27% | 0 | 0% | 6 | 40% |
| violence | 8 | 1% | 1 | 13% | 2 | 25% | 2 | 25% | 2 | 25% | 0 | 0% | 1 | 13% |
| visual features | 65 | 10% | 23* | 35% | 4 | 6% | 0 | 0% | 10 | 15% | 10* | 15% | 18*† | 28% |

*Indicates significant forward inference at *p*_corrected_ < 0.05 and †indicates significant reverse inference at *p*_corrected_ < 0.05 (corrected for false

### Automated Neurosynth Annotations

To complement the manual annotation analysis, we used Neurosynth’s automated annotations, which describes experiments that engage each MAG based on published neuroimaging data, allowing comparison of our corpus with the broader literature. MAG results were decoded in Neurosynth, yielding correlation values indicating the similarity of the input map (i.e., each MAG’s ALE map) and maps associated with each term from the Neurosynth database. To facilitate interpretation, the top ten terms with the highest correlation values for each MAG are presented (Table 5). Terms that were near-duplicates of terms already included in the list were removed, such as “*emotion*” and “*emotions*” if “*emotional*” was higher on the list. Non-content terms (e.g. “*abstract*”, “*reliable*”) and terms that described brain regions, such as “*insula*” or “*mt*”, were also excluded

**Table 5.** Automated functional decoding results from Neurosynth. The top ten Neurosynth (NS) terms are provided for each MAG, along with the corresponding Pearson’s correlation coefficient (“corr”) that indicates the strength of similarity between Neurosynth maps and each MAG

| MAG 1 |  | MAG 2 |  | MAG 3 |  | MAG 4 |  | MAG 5 |  | MAG 6 |  |
| --- | --- | --- | --- | --- | --- | --- | --- | --- | --- | --- | --- |
| NS term | corr. | NS term | corr. | NS term | corr. | NS term | corr. | NS term | corr. | NS term | corr. |
| motion | 0.555 | comprehension | 0.417 | neutral | 0.446 | navigation | 0.324 | sounds | 0.74 | visual | 0.431 |
| body | 0.451 | sentence | 0.408 | fearful | 0.437 | scenes | 0.316 | auditory | 0.732 | spatial | 0.414 |
| static | 0.441 | language | 0.375 | facial | 0.435 | episodic | 0.294 | listening | 0.711 | attention | 0.342 |
| moving | 0.415 | semantic | 0.351 | emotion | 0.434 | virtual | 0.278 | acoustic | 0.675 | eye movements | 0.300 |
| viewed | 0.406 | linguistic | 0.336 | expressions | 0.431 | memory | 0.276 | speech | 0.669 | execution | 0.299 |
| visual | 0.403 | theory mind | 0.318 | happy | 0.404 | retrieval | 0.270 | music | 0.625 | task | 0.286 |
| visual motion | 0.381 | mental state | 0.309 | angry | 0.401 | episodic memory | 0.258 | pitch | 0.612 | visuospatial | 0.279 |
| videos | 0.360 | mind | 0.306 | affective | 0.397 | place | 0.208 | spoken | 0.590 | movements | 0.274 |
| perception | 0.359 | mentalizing | 0.304 | facial expressions | 0.395 | autobiographical | 0.201 | tones | 0.572 | spatial attention | 0.256 |
| observation | 0.350 | language comprehension | 0.289 | neutral faces | 0.385 | remembering | 0.201 | voice | 0.568 | hand | 0.250 |

### Functional Interpretation of MAGs

The combined knowledge gained from the MAGs topography, as well as the manual and automated metadata decoding analyses provided insight into the functional interpretation of the clustering results. Overall, the terms yielded by Neurosynth decoding generally agreed with the manual annotation terms in characterizing the MAGs. Below is a summary of the six MAGs; note that reported labels do not refer to the definitive function of these regions, but rather indicate how each MAG reflects differential network contributions during naturalistic fMRI paradigms.

Manual annotations indicated that **MAG 1** experiments involved attention and the processing of dynamic visual features, in addition to visually-presented anthropomorphic forms and faces. Most of the stimuli in these experiments were films (Fig 4B), especially affective films. Neurosynth results largely converged with these manual annotations, as terms including “*videos”*, “*body*”, “*observation*”, and “*visual motion*” (Table 5) were associated with activations in MAG 1 regions. These annotations, together with the presence of convergent activation across regions commonly associated with higher-level visual processing, suggest that MAG 1 was associated with the ***Observation of Body and Biological Motion*** (Figure 3.1).

Manual annotations indicated that **MAG 2** experiments involved language processing, inference, and judgements about congruence. This MAG included relatively large proportions of the experiments using speech, video games, and tactile stimulation (Fig 3B). Neurosynth results supported the manual annotations’ indication that this MAG was associated with language processing and comprehension, as terms such as “*sentence*”, “*comprehension*”, “*semantic*”, and “*mentalizing*” (Table 5) were returned. These annotations and the presence of convergent activation in predominately left lateralized regions typically associated with higher-order cognition and language suggest that MAG 2 related to ***Language Processing*** (Figure 3.2).

Manual annotations indicated that **MAG 3** experiments involved human interactions or affective displays, including emotional and erotic films. Films were the predominantly used stimuli across these experiments, while most paradigms using painful stimuli were grouped into this MAG (Fig 4B). Neurosynth results corroborated these manual annotation interpretations regarding affective, aversive, and social processing, with terms such as “*emotion*”, “*facial expressions*”, “*fearful*”, and “*affective*” (Table 5). Together, these annotations and a convergent activation pattern involving bilateral amygdalae suggest that MAG 3 was associated with ***Emotional Processing*** (Figure 3.3).

Manual annotations indicated that **MAG 4** heavily represented experiments involving navigation through virtual reality environments, with spatial memory demands related to encoding unfamiliar virtual landscapes for future use. A few of these experiments required language processing, as well, and half of the experiments that used 3D images were grouped into MAG 5 (Fig. 4B). The manual annotations were reflected in the Neurosynth results, as similar patterns of activation have been associated with “*navigation*”, “*scenes*”, “*memory*”, and “*place*”. Additional related terms added depth to our characterization, expanding on the memory demands with “*retrieval*”, “*episodic memory*”, and “*remembering*” (Table 5). Overall, these experimental characteristics and convergent activation in medial temporal regions and along the visual processing stream suggest that MAG 5 was associated with ***Navigation and Spatial Memory*** (Figure 3.4).

Manual annotations showed that **MAG 5** experiments primarily involved either film or music stimuli (Figure 4B) and engaged either audiovisual or purely auditory processing (Fig. 4A). More than half of the included experiments that used music as stimuli were grouped into this MAG (Fig. 2B), with some stimuli involving an emotional quality (Table 4). Neurosynth corroborated these interpretations returning terms such as “*auditory*”, “*sounds*”, “*listening*”, and “*speech*” associated with activation of the regions in this MAG. These metadata descriptions combined with convergent activation in superior temporal regions suggest this MAG’s association with ***Auditory Processing*** (Figure 3.5).

Manual annotations of **MAG 6** experiments implicated tasks involving visual attentional demands and the processing of visual features, as participants engaged in video games, tactile stimulation, and virtual reality navigation (Figure 4B, Table 4). Stimuli with high visuospatial demands (i.e. video games, virtual reality, pictures, were represented more by this MAG than any other, while stimuli with low visuospatial demands (i.e. music, speech) were represented the least in this MAG. Some experiments involved memory encoding, and visual processing. Neurosynth supported this characterization returning terms including “*visual*,” “*attention*”, “*eye movements*”, “*saccades*”, and “*spatial attention*” associated with activation of the regions in this MAG (Table 5). These annotations and convergent activation in regions resembling the dorsal attention network and areas of higher level visual processing (e.g., superior frontal and parietal regions, extrastriate cortex) suggest this MAG’s association with ***Visuospatial Attention*** (Figure 3.6).

## Discussion

To characterize a core set of brain networks engaged in more ecologically valid neuroimaging designs, we employed a data-driven approach that meta-analytically grouped published naturalistic fMRI results according to their spatial topographies. Objective metrics suggested that a solution of *K* = 6 clusters provided the most stable and disparate grouping of experiments across the naturalistic fMRI literature, and ALE meta-analysis delineated convergent activation across spatially distinct brain regions for each meta-analytic grouping (MAG) of experiments. We then considered how such networks subdivide information processing by assessing the characteristics of the constituent experiments from each MAG. Utilizing both manual and automated functional decoding approaches, enhanced interpretations of the mental processes associated with specific constellations of brain regions were gleaned such that the outcomes of the two approaches generally agreed, with differences highlighting domain-specific and domain-general processes associated with naturalistic paradigms.

### Distributed Processing for Complex Functions

Though the six identified MAGs are spatially distinct and appear to correspond with dissociable mental processes, most of the included naturalistic tasks that reported more than one statistical contrast recruited more than one MAG (66 of 86). This is consistent with functional segregation and the flexible nature of the naturalistic design, demonstrating that the manipulation of different contrasts can identify distinct networks that likely cooperate to successfully perform a complex task. Further indicative of coordinated interactions and distributed processing, each MAG included experiments that utilized different task modalities and task types. Overwhelmingly, the identified MAGs and the functional characterizations thereof support the notion that complex behaviors are facilitated by coordinated interactions between several large-scale sensory, attentional, and domain-specific networks, a position increasingly endorsed in neuroimaging endeavors (Barrett and Satpute, 2013; Lindquist et al., 2012; Mišić and Sporns, 2016; Spreng et al., 2013). The characterization of identified MAGs from aspects of the naturalistic paradigms that elicit them suggest an information processing model of cooperating systems (Figure 5) for sensory input (MAGs 1 and 5), attentional control (MAG 6), and domain-specific processing (MAGs 2, 3, and 4), into and from which information is segregated and integrated to enable complex behaviors (e.g., language, emotion, spatial navigation). Output relevant to the corresponding input would be relegated by motor planning and execution systems, which are notable absent from the characterization of MAGs presented here, as experiments requiring a motor response were evenly distributed across MAGs, rather than clustered together.

**Figure 5.**
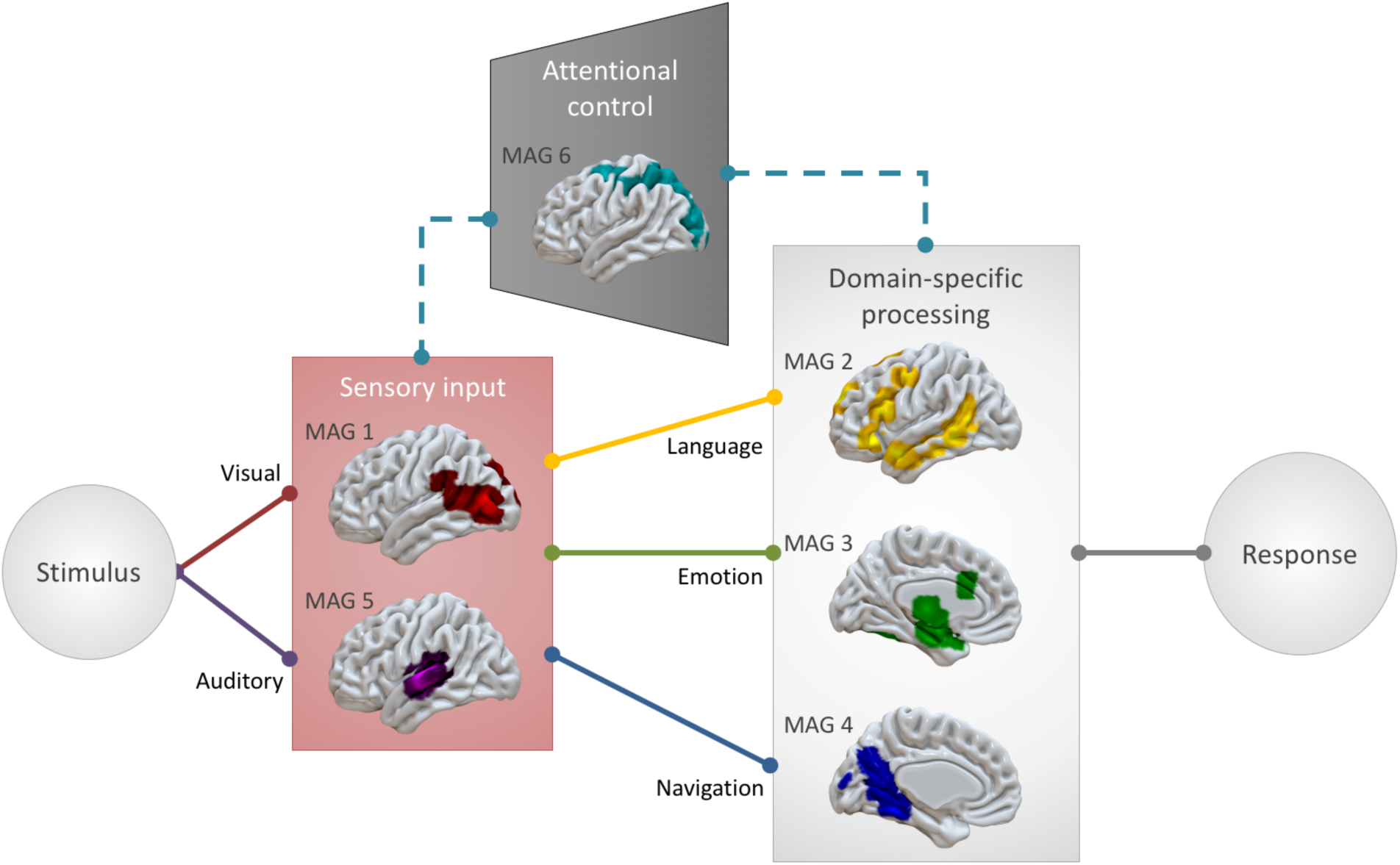
Complex systems for dynamical information processing. The identified MAGs present a framework of component systems that interact to enable complex information processing needed for naturalistic behavior, including necessary input systems, as well as systems for modality-specific (indicated by dashed line) visuospatial attentional gating of irrelevant information and domain-specific processing for language-, emotion-, and navigation-related tasks

MAGs 1 and 5 primarily represent the perceptual processing streams of incoming auditory and visual information, and likely cooperate to process audiovisual information. Functional decoding suggests that MAG 1 is involved in viewing faces and anthropomorphic figures, which is consistent with previous research showing that posterior temporal and temporo-occipital regions corresponding with area V5/MT are associated with the perception of movement, specifically biological movement (Cohen Kadosh et al., 2010; Pelphrey et al., 2005, 2004; Puce et al., 1998; Wheaton et al., 2004). Similarly, MAG 5 is associated with listening to music and speech, as well as perceiving pitch and tone, stretching across primary auditory cortex and into regions of higher auditory processing (Gray et al., 2009; Türe et al., 1999). Per functional decoding of MAG 6 of both manual and automated annotations, MAG 6 is associated with visuospatial attention. This functional characterization is also supported by corresponding fronto-parietal activations that are often associated with attending to visual stimuli (Braga et al., 2016; Puschmann et al., 2016). MAGs 1 and 5 represent the perceptual processing streams of audiovisual information.

Information processing depends on input from perceptual systems, filtered by attentional gating, but proceeds in a functionally-segregated manner, seen in domain-specific MAGs for linguistic, emotional, and spatial processing. When considering language processing, there is necessary input to primary auditory areas (MAG 5), that is further processed by higher-level language areas that facilitate speech perception and comprehension (MAG 2). More than a third of contrasts from experiments that utilized speech-based paradigms contributed to the convergent activation pattern of MAG 2, which was linked by both functional decoding techniques to language-related processes. Furthermore, the regions of MAG 2 resembles a neural “language network” (Friederici and Gierhan, 2013; Heim et al., 2003; Price, 2010; Saur et al., 2010), including some regions associated with orofacial articulation (lip, tongue, and jaw movements) and motor planning (SMA, pre-SMA) that allow the motor components of speech. By presenting language in a context that is more representative of how we process language in everyday life, such as through the use of spoken fictional narratives (AbdulSabur et al., 2014; Wallentin et al., 2011; Xu et al., 2005a) or scene descriptions (Summerfield et al., 2010), naturalistic fMRI paradigms allow researchers to explore the multiple neural networks at work in performing the cooperating processes that facilitate language processing. Similarly, emotional processing (MAG 3) often necessitates audiovisual input (MAGs 1 and 5) and necessitates attention (MAG 6). Emotional films recruited regions across these four MAGs, suggesting a similarly diverse group of coordinated neural systems are engaged when observing affective displays. Additionally, navigation (Burgess et al., 2002; Kalpouzos et al., 2010; Wolbers et al., 2004) depends on visual input (MAG 1), effective visuospatial attentional (MAG 6), and spatial memory and processing (MAG 4). The functional characterization of MAG 4 from manual and Neurosynth decoding highlights its involvement in navigation and spatial memory, supported by studies of rats and humans with brain lesions that indicate the importance of medial temporal, hippocampal, and precuneus regions in processing visual scenes and spatial information (Bird and Burgess, 2008; Epstein, 2008; Lee et al., 2005; Sailer et al., 2000; Squire et al., 2004; Summerfield et al., 2010; Xu et al., 2005b).

Finally, the characterization of MAG 6 indicates a domain-specific attentional system, as both manual and automated Neurosynth decoding highlight its involvement in visual processing in the absence of any association with other modalities. This is reflected by the distributions of stimuli across MAGs (Figure 4), which show low numbers of auditory and pain-related stimuli represented in MAG 6, while rich visual stimuli that include spatial information (i.e. video games, virtual reality, pictures) are highly represented across the experiments in MAG 6. Curiously, tactile object manipulation was highly represented in MAG 6, representing the perception of spatial information in the absence of visual information (Figure 4, Table 4). Together, these suggest that MAG 6 provides modality-specific attentional gating, depicted by the dashed line in Figure 5.

### Limitations

The present results may be limited by the *k*-means clustering method, which is limited by the assumptions of the algorithm and underlying topology of the data, as it is sensitive to spherical clusters and assumes the data are linearly separable. Furthermore, there is a potential for bias with this method, certain parameters are specified by the researcher beforehand. To address this potential for bias and the stability of our clustering solution, we performed duplicate clustering analyses with both linear (hierarchical clustering using Ward’s method) and nonlinear (kernel *k*-means and density-based spatial clustering) methods. The results of these analyses are provided in the Supplementary Material (Supplementary Figures 1 – 3) and confirmed that our choice of the *k*-means clustering method provided optimal separation of the data into 6 clusters. Experiments in our corpus were grouped using the *kmeans++* algorithm for each of *K* = 2 through *K* = 20 solutions, repeated 1000 times to ensure that each solution minimized the point-to-centroid distance, indicative of optimal clustering (Kanungo et al., 2004). Pearson’s correlation was selected as the distance metric, as recommended by Laird et al. (2015). The *K* = 6 solution was designated as an optimal candidate solution before assessing the convergent activation patterns of each MAG, based on the aforementioned metrics, yielding a data-driven result. These results are, of course, influenced by the choice of clustering method, and should be considered accordingly. As this was a meta-analytic effort, it is limited, too, by the initial modeling of the data. Despite this, coordinate-based meta-analyses are considered a robust method for synthesis of previously published functional neuroimaging literature (Eickhoff et al., 2012, 2009; Fox et al., 2005). Although the functional decoding based manual annotations relied on a subjective process, the results were largely confirmed by comparison with the wider body of functional neuroimaging literature facilitated by Neurosynth’s automated functional decoding. It is worth noting that the naturalistic literature is somewhat limited, with an emphasis on navigation and affective processing, and continued research and expansion of this corpus will facilitate development of a more comprehensive model of the neural networks that support realistic behavior.

### Summary and Future Work

In summary, this meta-analysis of naturalistic fMRI studies that apply dynamic, lifelike tasks to explore the neural correlates of behavior has shown that these paradigms engage a set of core neural networks, supporting both separate processing of different streams of information and the integration of related information to enable flexible cognition and complex behavior. We identified seven patterns of consistent activation that correspond with neural networks that are involved in sensory input, top-down attentional control, domain-specific processing, and motor planning, representing the set of behavioral processes elicited by naturalistic paradigms in our corpus. Across the corpus, tasks provided mainly visual and auditory sensory input which engaged regions across MAGs 1 and 5, while MAG 6 appeared to contribute to top-down attentional control to filter out nonessential visual and/or spatial information. Salient information can be processed by the relevant domain-specific networks, shown in MAGs 2 (language), 3 (emotion), and 4 (navigation and spatial memory), informing the appropriate response. Most naturalistic tasks engaged multiple networks to process the relevant information from a stimulus and generate an appropriate response. A shift in favor of utilizing naturalistic paradigms, when possible, would greatly benefit the field, as naturalistic stimuli more closely approximate the full complement of processing necessary for realistic behavior. Due to the availability of naturalistic fMRI data from sources such as studyforrest.org, the Human Connectome Project, and the Healthy Brain Network Serial Scanning Initiative (HBNSSI), an intriguing next step in this line of work would include validating these MAGs in the primary analysis of imaging data. Exploring how multifaceted processes interact and, ultimately, contribute to behavior will allow us to better understand the brain and human behavior in the real world. In the future, studies of this sort would greatly benefit from an automated annotation process for an objective functional decoding of included papers, instead of subjective manual annotation.

## Accessibility of Data and Other Materials

The authors have released all code and data associated with this manuscript. The code and tabular data are available on GitHub (https://github.com/62442katieb/meta-analytic-kmeans), and the unthresholded maps of each MAG are available on NeuroVault (https://neurovault.org/collections/3179/).

## ACKNOWLEDGMENTS

This study was supported by awards from the National Institute of Drug Abuse (U01-DA041156, K01-DA037819, U24-DA039832, R01DA041353), the National Institute of Mental Health (R56-MH097870), and the National Science Foundation (1631325 and REAL DRL-1420627). The authors declare no competing financial interests.

